# Disease and ectoparasite management improve nestling golden eagle health and survival: An effective mitigation strategy

**DOI:** 10.1101/2024.08.14.607861

**Authors:** Julie A. Heath, Caitlin M. Davis, Benjamin M. Dudek, Christopher J. W. McClure, Kevin T. Myers, Emma K. Regnier, Brian W. Rolek, Ashley L. Santiago

**Author notes:** **Corresponding Author**: Julie Heath. **Open Research statement:** Data and code are available at an institutional DOI (TBD).

## Abstract

Changes in the distribution and behavior of pathogens and parasites associated with climate change are creating emerging threats for wildlife. These threats have created an urgent need for conservation strategies to manage wildlife. Conservation strategies could provide reliable and cost-effective compensatory mitigation options for protected species affected by renewable energy production, such as golden eagles (*Aquila chrysaetos*). We used nest camera data to assess survival and cause of death for golden eagle nestlings in southern Idaho, USA. Together, poultry bugs (*Haematosiphon inodorus*), a blood-sucking ectoparasite that lives in golden eagle nests, and trichomonosis, a fatal disease that is transmitted through the consumption of rock pigeons (*Columba livia*) caused ∼ 27% of nestlings to die per year. From 2022—2023, we treated trichomonosis-infected nestlings with an anti-protozoan drug and developed a protocol for reducing poultry bug densities in eagle nests, then we projected the effects of disease and parasite treatments on population growth using simulation models. We collected pre-treatment information on nestling health and survival, and then, after young fledged, we applied 1 of 3 experimental parasite treatments to eagle nests: permethrin and diatomaceous earth, diatomaceous earth only, or water – a control, and measured effects on eagle health and survival the following year. Ten eagle nestlings with trichomonosis, that otherwise would have died, were successfully treated and lived to fledge from the nest. The permethrin and diatomaceous earth treatment significantly reduced poultry bug densities in eagle nests in the subsequent breeding season and created measurable improvements to nestling health and survival. Nestlings had significantly higher hematocrit, female nestlings had higher mass, and, on average, more nestlings (0.70) successfully survived to fledge compared with control- and diatomaceous earth-treated nests. Further, treatment of disease and parasites was projected to increase population growth rates by 4% and 8%, respectively. In total, we estimate saving 17.5 nestlings, which equates to 10.2 adult eagles, through disease and parasite treatments in two years. In areas with dense concentrations of poultry bugs or where eagles consume rock pigeons, disease and parasite treatments could be an effective compensatory mitigation option or management tool with population-level impacts on eagle populations.

## Introduction

Climate change is altering the distribution and behavior of various pathogens and parasites, leading to an increased risk for wildlife (Harvell et al., 2002). For example, rising temperatures and changing precipitation patterns can expand the geographical range of disease vectors, exposing wildlife populations to novel infections (Lafferty, 2009), or increase the overwinter survival of parasites or vectors, leading to greater infection risk (Bale and Hayward, 2010, although see Brown et al., 2022). Additionally, landscape changes might disrupt ecosystem dynamics, making some species more susceptible to infections (Hassell et al., 2013). The compounding effects of environmental change on disease dynamics underscore the urgency of comprehensive conservation strategies to mitigate these threats to wildlife health, particularly for species of conservation concern (Smith et al., 2006).

Apex avian predators (raptors) are declining worldwide (McClure et al., 2018) and they are particularly susceptible to parasites and pathogens (Blanco et al., 2022; van den Brand et al., 2015; Vidaña et al., 2020). Raptors often migrate long distances or have large home ranges, which can increase exposure to pathogens (Altizer et al., 2011). Further, raptor nests are large and provide optimal ectoparasite habitat (Philips and Dindal, 1977) and an easily accessible food source (i.e., nestlings) that cannot leave the nest until fledging (Philips, 2007). Finally, ecosystem changes in trophic systems can expose raptors to disease through novel interactions with pathogens or vectors. For example, as the preferred prey of Bonelli’s eagles (*Aquila fasciata*) decreases, eagles switch prey to forage on rock pigeons (*Columba livia*) that carry the protozoan *Trichomonas gallinae,* which can cause the disease trichomonosis and nestling death (Palma et al., 2006; Real et al., 2000).

Golden eagles (*Aquila chrysaetos*) in western North America are affected by disease and parasites. Like Bonelli’s eagles, when faced with habitat degradation and reduced availability of preferred prey such as black-tailed jackrabbits (*Lepus californicus*), golden eagles switch to alternative prey such as rock pigeons the vectors of *T. gallinae* that cause trichomonosis (Heath et al., 2021). Trichomonosis is characterized by plaques in the oral cavity that grow to block the esophagus and trachea of nestling eagles, resulting in a high mortality rate (Dudek et al., 2018). Further, poultry bugs (*Haematosiphon inodorus*, also known as Mexican chicken bugs), are a blood-sucking ectoparasite in the family Cimicidae and are closely related to bed bugs (*Cimex* spp.) and swallow bugs (*Oeciacus vicarius*). The effects of poultry bugs on golden eagles have been documented for several decades in the southwestern US (Grubb et al., 1986; Wilson and Oliver, 1978). More recently, the incidence and negative effects of poultry bugs on golden eagle populations have increased (Murphy et al., 2023), perhaps because warming winters (Møller et al., 2013) have led to the rapid expansion of the poultry bugs’ range (McFadzen et al., 1996). Consequently, poultry bugs might be considered an emerging threat to golden eagles in much of their western range. Poultry bugs live in nest material and emerge to feed on the blood of nestlings. Poultry bugs reduce golden eagle nestling mass, blood hematocrit, and telomere length (all correlates of nesting health), increase corticosterone (a hormone associated with physiologic stress), and increase the probability that nestlings fledge early (and die) or die in the nest (Dudek et al., 2021). Young eagles that successfully survive to fledge from a nest with poultry bugs have a low probability of survival in their first year (Murphy et al., 2023).

Treatments for pathogens and parasites vary in approach and efficacy. Trichomonosis can be treated if the infection is discovered early when the plaques are small. One dose of an anti-protozoan reverses the infection and nestlings survive. Unfortunately, effective treatment options for poultry bugs remain unclear. Organically derived products such as diatomaceous earth (Bennett et al., 2011; Dawson, 2004; Kochert pers comm.) and pyrethrins (Fessl et al., 2006; Hanssen et al., 2013) have previously been used to treat ectoparasites, including some from the family Cimicidae (swallow bugs and poultry bugs). Diatomaceous earth abrades the cuticle of the parasite, causing it to desiccate (Korunić, 2013). Pyrethrins and the synthetic version, permethrin, impact the nervous system by disrupting the sodium ion channel and paralyzing the insect (Soderlund et al., 2002). Though widely used to treat nest parasites, pyrethroids applications can have side effects on young birds (Larsen et al., 2004, López-Arrabé et al., 2014) or their offspring (Bulgarella et al., 2020) and may contain dangerous additives (Hund et al., 2015). The most effective timing and dosages for the use of these products as a treatment for poultry bugs in eagle nests, while minimizing impacts to nestling eagles, remains uncertain.

Given the increasing threat of disease and parasites, and the consistent timing (breeding season) and location (nest) of their effect, the treatment of disease and parasites may be an excellent management tool or mitigation option. Several US laws, including the Bald and Golden Eagle Protection Act (16 U.S.C. 669-668c), protect eagles by preventing ‘take’, the killing, wounding, or harassment of birds. The United States Fish and Wildlife Service manages the permitting for individuals or organizations that may incidentally or unavoidably take eagles while conducting otherwise lawful activities. As part of the permitting process, permittees are often required to offset their take through mitigation. The US has a growing dependence on large-scale renewable energy; particularly wind power. Mortality of juvenile and adult golden eagles due to collisions with wind turbines has been documented (Millsap et al., 2022), creating a need to identify reliable direct or compensatory mitigation options. Unfortunately, few mitigation techniques can quantifiably demonstrate a net positive impact on eagle populations (Allison et al., 2017). Increasing the number of mitigation options will benefit renewable energy permittees and eagle populations by focusing conservation efforts on removing sources of eagle mortality. To date, the reduction of eagle mortality caused by nest ectoparasites or disease has not been widely considered for offsetting incidental eagle take.

Our objective was to examine the effects of disease and parasites on golden eagle nestlings in southern Idaho, USA where golden eagles reach some of the highest nesting densities in North America. Further, we aimed to treat nestlings for trichomonosis and develop an effective treatment option for poultry bugs that minimizes potential side effects to eagles. The incidence of trichomonosis in golden eagle nestlings has increased in southern Idaho with ∼ 41% of nestlings exposed to *T. gallinae* and 25% of nestlings developing lethal oral plaques (Dudek et al., 2018). In the same study area, poultry bugs were reported in golden eagle nests as early as the late 1960s (Hickman, 1968). More recently, Dudek et al. (2021) found the presence of poultry bugs in golden eagle nests to be widespread, affecting 77% of eagle nests. The consistent and high rate of poultry bug infection of nests in southern Idaho provides an excellent opportunity to test the efficacy of anti-parasite treatments on poultry bug presence and abundance and nestling health and survival. We aimed to assess the efficacy of poultry bug treatment strategies by testing two readily available anti-parasite products (diatomaceous earth and permethrin) on nests of golden eagles in southern Idaho. Then, we projected whether trichomonosis or poultry bug treatment could have population-level effects. Finally, we considered whether nestling parasite and disease treatment may be a viable compensatory mitigation option for golden eagle incidental take permittees.

## Methods

Our study area was located in southwestern Idaho along the Snake River Canyon in the Morley Nelson Snake River Birds of Prey National Conservation Area and surrounding areas (Figure 1). Steep basalt cliffs in the Snake River Canyon and rocky outcrops provide ample sites for golden eagles to construct large stick nests. The surrounding uplands consist of native shrub-steppe and salt-desert communities characterized by sagebrush-dominated shrublands, native perennial grasses, disturbed grasslands, rangeland dominated by exotic annual grasses, irrigated agricultural land, and rural and suburban development (U.S. Department of the Interior, 1996).

**Figure 1.**
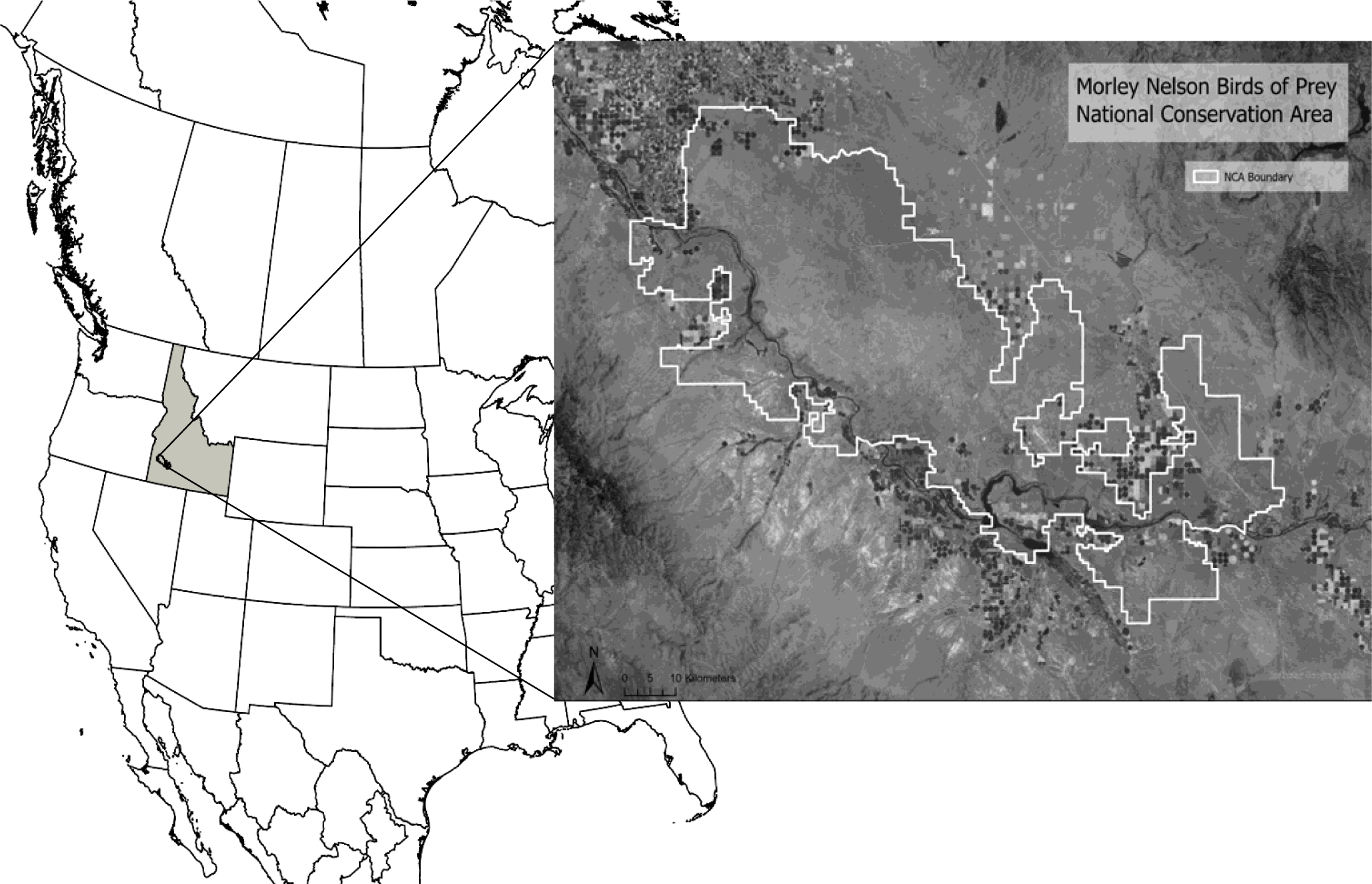
Map of the study area in the Morley Nelson Snake River Birds of Prey National Conservation Area (NCA) in southern Idaho, USA.

### Effects of disease and parasites on nestling health

We monitored 80 – 90 historical golden eagle nesting territories in southern Idaho in 2014 – 2015, 2017 – 2018, and 2021. We followed ground-based survey methods described by Steenhof et al. (1997) to determine territory occupancy and nesting attempts. Once we identified a nesting attempt, we made regular nest observations from >400 meters to monitor outcomes and visually assess nestling age based on feather development (Driscoll, 2010). We rappelled into 15 – 25 nests per year to install trail cameras (Harrison et al., 2019) when nestlings were 2 – 3 weeks old and able to thermoregulate (Katzner et al., 2020). Trail cameras were set to record images with either a motion-activated trigger or a time-lapse setting of 1 image per hour. Once we did not observe young in the nest, we returned to collect the cameras and review images. We used camera images to determine the date and age that nestlings fledged (left the nest) or died.

The cause of death was based on images and information we collected while visiting the nest. When we installed cameras, we examined nestling eagles for trichomonosis plaques by visually inspecting the sides of the mouth, oropharynx, and upper and lower surfaces of the tongue. If we detected a possible lesion, we used a cotton swab to probe the lesion to test whether it was attached or mobile (e.g., a piece of food). We considered immobile, caseous lesions to be indicative of trichomonosis. If nestlings had trichomonosis, we treated individuals with anti-protozoan medication (see below). Additionally, during our nest visit, we recorded whether there were poultry bugs in the nest. We considered the cause of death to be trichomonosis if a nestling had plaques in its mouth (treated or not). We considered the cause of death to be poultry bugs if a nest was infested and a nestling died of starvation (despite prey deliveries), or if a nestling jumped from an infested nest to its death before it could fly.

### Disease and Parasite Treatment

Our study of disease and parasite treatments occurred in the breeding seasons of 2021 – 2023. During all visits to nests with nestling eagles, we examined nestling eagles for trichomonosis plaques as described above. Nestlings with caseous lesions were treated with a 30 mg oral dose of carnidazole (Spartrix, Harkers, Suffolk, UK), an antiprotozoal drug, to reduce the growth of plaques that would otherwise lead to nestling death. The carnidazole was administered orally, by placing the tablet in the back of the esophagus so it was swallowed. We followed the fate of these nestlings with camera images (see below).

Before parasite treatments, in 2021, we selected 18 occupied eagle territories with nesting attempts for our pre-treatment assessment of poultry bugs. We visited nests when eagle nestlings were ∼ 3 weeks old, to bury 2 – 3 SenSci ActivVolcano Bed Bug Detector® (SenSci, Lawrenceville, NJ, USA) traps within eagle nests. These small pitfall traps had a scent lure designed to attract (from < 1 m) and capture common bed bugs (*Cimex lectularius*), a closely related species to poultry bugs. Traps were covered with a small cardboard ‘tent’ to prevent nest material from falling into the trap. We took photographs and mapped the locations of the traps to aid in the recovery of traps during the next visit. We returned to the nests again ∼ 2 months later, once the nests were empty, to retrieve the traps (and apply treatments, see below). We placed recovered traps in sealable plastic bags and stored them in the freezer until the end of the season when we tabulated trap contents.

We randomly assigned an equal number of territories to one of three treatment options: 1) diatomaceous earth, 2) a combination of diatomaceous earth and permethrin (Permacap, BASF, NJ, USA), or 3) a water-only (control) treatment. We considered eagle territories to be the experimental units for our study of parasite treatments. Golden eagle nesting territories often contain multiple alternative nests (Millsap et al., 2015). In southern Idaho, the number of nests per territory can range from 1 to 18 with some pairs using the same nest every year, whereas other pairs switch nests within their territory from year to year (Kochert and Steenhof, 2012). We aimed to treat the three most recently used, and climber-accessible, nests within a territory. By selecting multiple, recently used nests we increased the probability that the eagle pair would use the treated nest in the subsequent breeding season (see below, Kochert and Steenhof, 2012; Wiens et al., 2018). Further, adult eagles regularly perch on alternate nests and may act as vectors by carrying poultry bugs from one nest to another. By treating several nests, we aimed to decrease territory-level poultry bug load.

We applied all parasite treatments in the summer (mid-June – July) after the young had fledged. By waiting until the young left the nest, we decreased researcher disturbance and avoided potential secondary effects on nestling eagles from the treatments (López-Arrabé et al., 2014). Despite nestlings eagles being out of the nest, the poultry bugs reside in eagle nests year-round and remain active through the fall, leaving them vulnerable to treatment (Heath unpub. data). We anticipated that a summer treatment would decrease annual poultry bug survival, leading to a lower density of poultry bugs in eagle nests in subsequent springs. Diatomaceous earth powder was dusted over the surface of the nest continuously, resulting in approximately 60 grams of treatment product being applied per nest (Dawson, 2004; Hanssen et al., 2013). Fluid-based treatments (permethrin and water) were applied via a hand-pumped garden sprayer at a rate of ∼ 290 ml per square meter. Permacap was diluted to 1:10 before application. The crew wore personal protective equipment including gloves, eye protection, and a respirator, when applying all treatments. The treatment applied to each territory was randomly selected and usually varied between years. Six territories were treated in only a single year, and 18 territories were treated in both years, though not always with the same treatment. In some cases, treated territories were not occupied in subsequent years, pairs did not attempt to breed, or nests were inaccessible for climbing. At these territories, we were not able to collect post-treatment data on either parasite load in the nest, nestling health, or both.

In 2022 and 2023, we monitored focal eagle territories in late winter and early spring (Jan – Mar) for occupancy and observation of incubating eagles. We aimed to visit treated territories when the young were approximately 21 days old and again when they were 45 – 56 days old (6.5 – 8 weeks). On the first nest visit, we placed SenSci ActivVolcano Bed Bug Detector® traps in nests as described above and installed a nest camera to record nestling fate. On the second visit, we recorded morphometric measurements, including mass (a measure of condition) and footpad length (a size measure that can be used to determine nestling sex), and used a 25-gauge needle and syringe to withdraw < 1 ml blood from the brachial vein to measure nestling hematocrit. Finally, we returned to the nests to collect the traps and camera once all the young in the nest were gone. We reviewed images from nest cameras to determine whether or not nestlings successfully survived to fledge.

### Data synthesis and analysis

We estimated the effects of treatment for trichomonosis using an intercept-only Poisson generalized linear mixed model (GLMM) with the number of nestlings surviving to fledge per nest (hereafter nestling survivorship, Heath et al. 2021) as the response variable. This model included data from all focal nests, even where trichomonosis was not detected (i.e., treatment survivorship = 0), to estimate the effects of treatment study-wide. The result is thus the number of fledglings produced by the treatment per visited nest. We included territory identity as a random effect because some territories were sampled in more than one year. We considered the effect to be statistically significant if the intercept was significantly different from zero. We estimated “control” survivorship for the trichomonosis treatment using the same model except that the dependent variable was the number of offspring that would have fledged without treatment, i.e., the actual number that fledged minus the number treated. In both models, the year and the number of visits per territory did not affect the number of young that survived to fledging and these terms were removed.

We tallied all of the poultry bugs collected per nest. We used a GLMM with a negative binomial distribution to examine whether period (pre- or post-treatment), treatment, or the interaction between period and treatment explained poultry bug counts. This model included an offset for the number of traps retrieved from each nest and a random effect of territory identity because some territories were sampled in more than one year. We used general linear mixed models to examine whether treatment affected nestling hematocrit or mass. We used sex as a covariate for the mass model because female eagles are larger than male eagles. Both morphometric models had a random effect for each nest event identity to account for data collected from siblings.

We used a GLMM with a Poisson distribution to examine the effect of parasite treatment on nestling survivorship. This model included a fixed effect of parasite treatment type and random effects of year (categorical) and territory identity. For all analyses, we used a one-tailed null hypothesis, with the alternative hypothesis that more young survived to fledge in treated nests compared to controls. The one-tailed approach was used because we predicted that parasite and disease treatment would only increase nestling survivorship and health and to decrease the probability of a type II error given the relatively small sample sizes and possible small effect sizes (i.e., eagle broods range from 0 – 3 young).

### Population Effects

To inform potential mitigation strategies involving the mortality of golden eagles from wind turbine collisions, we used previously published rates of survival (Millsap et al. 2022) and our nest survivorship estimates with simulations to project the relative number of adult golden eagles and population growth rates produced by rescuing nestlings from disease and parasites. We used nestling survivorship as a proxy for fecundity, though this estimate is biased high because it does not include the fates of breeding attempts that failed early (i.e., < 3-week-old nestling), before we were able to install a camera in the nest. However, the nestling survivorship estimates are reasonable for comparing the relative effect size for disease and parasite treatments – effects that occurred once young were older than 3 weeks. We estimated nestling survivorship with and without disease and parasite treatments using the Poisson GLMMs described above. These models produced mean estimates of nestling survivorship on the log scale and their standard errors for treatment and control groups. We estimated fecundity, the number of female young produced per female, as nestling survivorship divided by 2 thereby assuming an approximately equal sex ratio.

We used simulation to examine the effects of our disease and parasite treatments on the projected population growth rate (Lambda, λ). We implemented simulations using the four-stage Lefkovitch matrix from Millsap et al (2022) to examine population dynamics of golden eagles across North America. We parameterized this matrix using survival rates estimated by Millsap et al. (2022) and fecundity rates either with or without disease treatment or parasite treatment. We modified code provided by Schaub and Kéry (2022; page 85) to randomly sample 10,000 survival values from a beta distribution, given average values of survival and their standard errors.

Using model-estimated values of fecundity, we randomly drew 10,000 times from a normal distribution as the mean and standard deviation on the log scale. Next, we exponentiated these values to backtransform and divided them by two to obtain random draws of fecundity (i.e., female only). We first projected the effects of trichomonosis treatment on λ. We used the mean and standard error of the control level of fecundity on the log scale outputted from the GLMM described above and randomly drew 10,000 times from a normal distribution. Then, to estimate the level of fecundity under the treatment regime, we added 10,000 draws of the effect size to the draws of the control level of fecundity. We exponentiated these values and divided them by two to obtain random draws of fecundity and parameterized 20,000 Lefkovitch matrices using these draws of fecundity either with or without treatment and Millsap et al.’s (2022) survival estimates. We performed a similar procedure to project the effects of treatment for poultry bugs on λ. We again parameterized 20,000 Lefkovitch matrices using these draws of fecundity either with (10,000 draws) or without (10,000 draws) parasite treatment, along with the random draws of survival mentioned above.

We used these random draws of vital rates to project the number of adult golden eagles resulting from our treatments. For each set of draws, we calculated the increase in fecundity because of disease or parasite treatment then multiplied it by two to calculate the number of offspring fledged (male and female). We then multiplied this value by each of the draws of first-, second-, and third-year survival. This produced the number of adult eagles (age ≥ 3) resulting from each territory treated. To estimate the number of adult eagles resulting from trichomonosis treatment, we multiplied this value by the number of territories checked for trichomonosis (27). To project the number of adult eagles resulting from parasite treatment, we multiplied the number of adult eagles resulting per territory by the number of territories treated for parasites (10). Our simulations yielded 10,000 values of λ and the number of adults resulting from treatment. We added these draws of number of adult eagles together to determine the number of adult eagles resulting from both treatments. We used the means, 5^th^, and 95^th^ percentiles to describe the central tendency and precision of these distributions.

## Results

### Effects of disease and parasites on nestlings

We installed cameras in 56 nests across 33 territories from 2014 – 2015, 2017 – 2018, and 2021. We monitored the fate of 87 nestlings in these 56 nests. Thirty-four of 87 nestlings (39%) died between 3 weeks old and fledging. The most common causes of death were poultry bugs (35%), and trichomonosis (35%). In some cases, nestlings had both poultry bugs and trichomonosis (6%). In total, 76% of deaths were caused by parasites, disease, or both. There were some unknown causes of death (n = 6, 18%), one nestling died from siblicide, and another of infanticide (6% combined).

### Disease and Parasite Treatments

We examined 69 nestlings (2022 n = 28, 2023 n = 41) at 27 nests for trichomonosis and detected plaques in 12 nestlings (2022 n = 5, 2023 n = 7). We successfully treated 10 eagle nestlings for trichomonosis and these individuals fledged. In two other cases, we discovered the disease at a late stage, when the birds had large plaques. Although we treated these birds, they did not survive because the disease was too progressed. On average, nestling survivorship significantly increased by 0.37 nestlings per nest (90% CI: 0.22 **—** 0.62) because of trichomonosis treatment.

We treated 14 territories (34 nests) with permethrin and diatomaceous earth (2021 n = 7, 2022 n = 7), 13 territories (35 nests) with diatomaceous earth (2021 n = 7, 2022 n = 6), and 15 territories (40 nests) served as controls (2021 n = 8, 2022 n = 7). In some nests, adult eagles turned poultry bug pitfall traps over or removed them from the nest and they could not be sampled. Therefore, our post-treatment sample size for poultry bug traps included 8 permethrin and diatomaceous earth territories, 4 diatomaceous earth territories, and 6 control territories. Poultry bug loads were similar across all groups in the pre-treatment period. We found a significant interaction between treatment type and period on poultry bug loads. Permethrin and diatomaceous earth treatment significantly decreased the poultry bug load in the post-treatment period (β_pre*permethrin and diatomaceous earth treatment_ = 5.1 ± 1.6, z = 3.2, p < 0.01, Figure 2), but there was no effect of treatment on post-treatment poultry bug loads in the diatomaceous earth (β_pre* diatomaceous earth treatment_ = 1.7 ± 1.7, z = 1.0, p = 0.16, Figure 2) or control groups (β_pre* control_ = 0.4 ± 0.8, z = 0.5, p = 0.31, Figure 2). Specifically, there were no highly infested nests in the permethrin and diatomaceous earth treatment group, which greatly lowered the variance (Figure 2A) and the overall mean (Figure 2B) poultry bug load. We combined the diatomaceous earth and control treatment groups into the control group for comparisons to the permethrin and diatomaceous earth treatment in analyses of nestling health and survival because there were no significant differences between the diatomaceous earth and control groups on poultry bug load.

**Figure 2.**
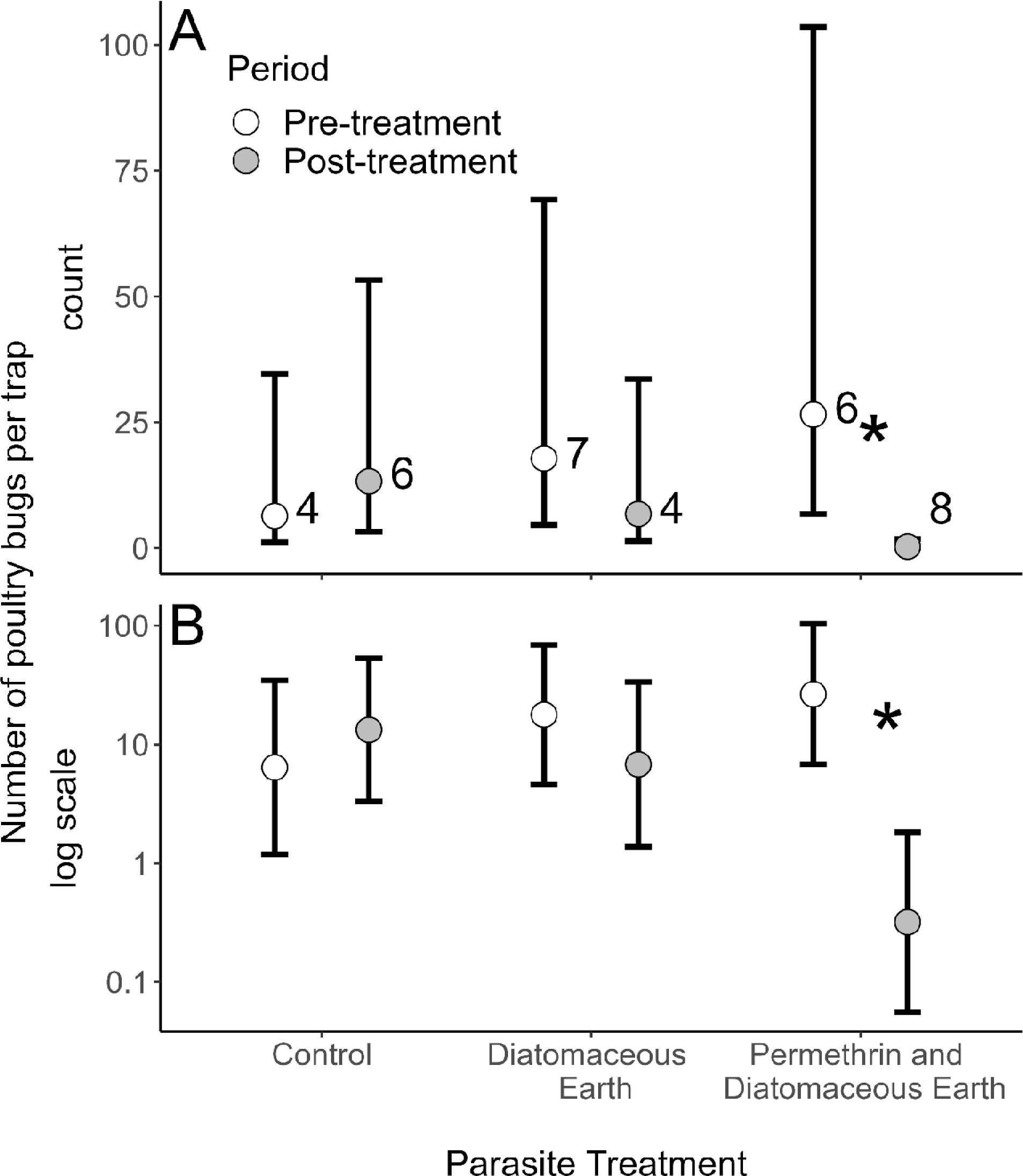
The effects of parasite treatments on the number of poultry bugs per trap (A, count and B, log scale) in golden eagle nests in southern Idaho, 2021-2023. There was a significant decline in parasite load in the permethrin and diatomaceous earth group after treatment. Numbers represent sample sizes; circles represent means and bars represent 90% confidence limits.

We found that eagle nestlings in the permethrin and diatomaceous earth treatment group had significantly higher hematocrit levels than nestling eagles from the control group (β_treatment_ = 5.8 ± 2.7, χ^2^ = 4.7, p = 0.02, Figure 3). Further, female nestling eagles had significantly higher mass than female nestlings from the control group (β_treatment_ = 686 ± 254, t = 2.7, p = 0.03, Figure 4). There was no effect of treatment on male mass (β_treatment_ = 178 ± 214, t = 0.83, p = 0.42, Figure 4). Finally, more nestlings successfully fledged from nests that had been treated with permethrin and diatomaceous earth compared to the control group (β_treatment_ = 0.76 ± 0.4, χ^2^ =3.4, p = 0.03, Figure 5). On average, 0.70 (90% CI: 0.18 — 1.22) more nestlings successfully fledged from nests in permethrin and diatomaceous earth-treated territories (1.30 nestlings per nest, 90% CI: 0.82 — 2.05) compared with control group territories (0.61 nestlings per nest, 90% CI: 0.37 — 1.00).

**Figure 3.**
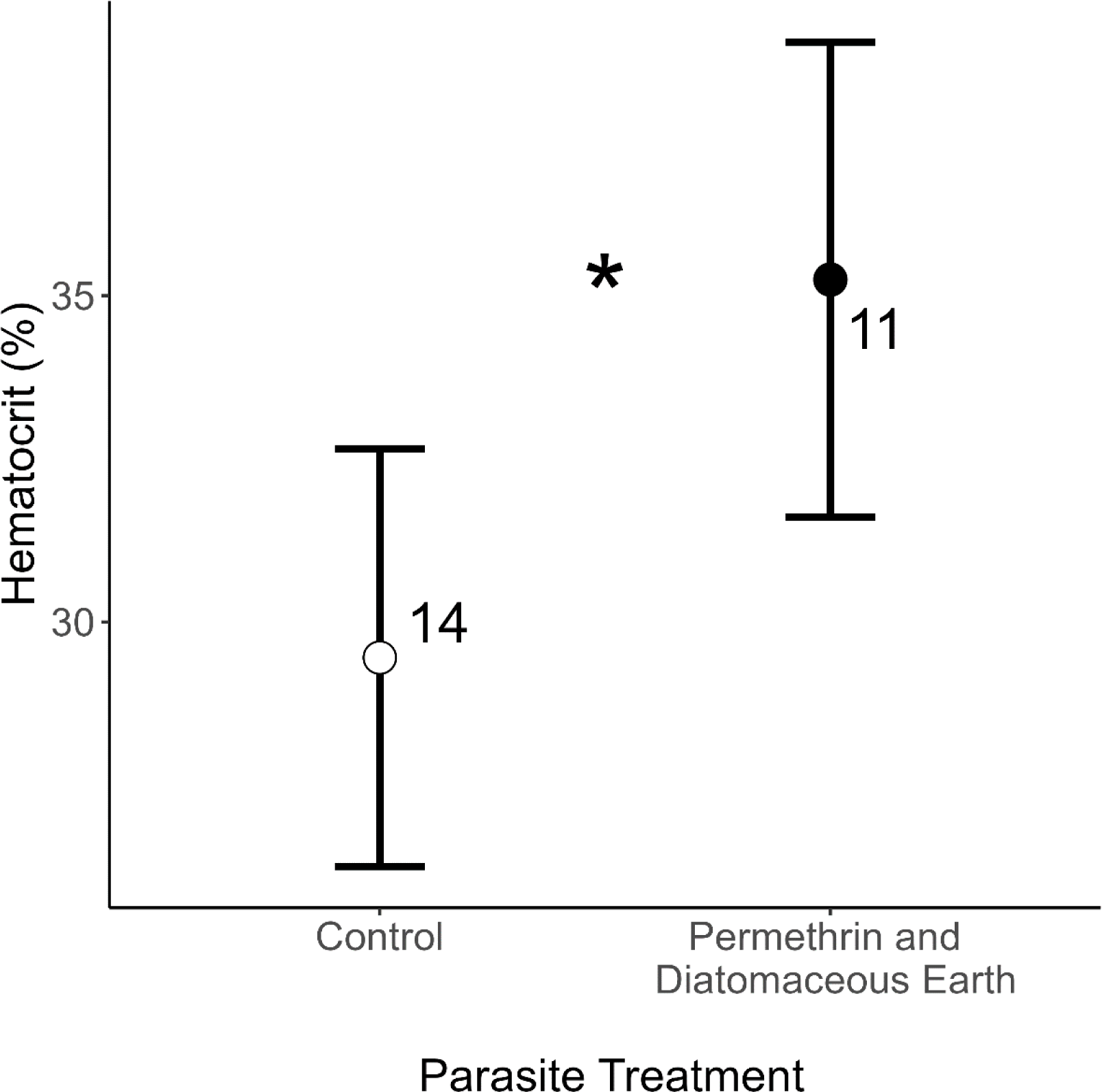
The effect of parasite treatments on the hematocrit of golden eagle nestlings > 5 weeks old. Nestlings in control nests, which had high poultry bug loads, had significantly lower hematocrit than nestlings in treated nests. Numbers represent sample sizes; circles represent means and bars are 90% confidence limits.

**Figure 4.**
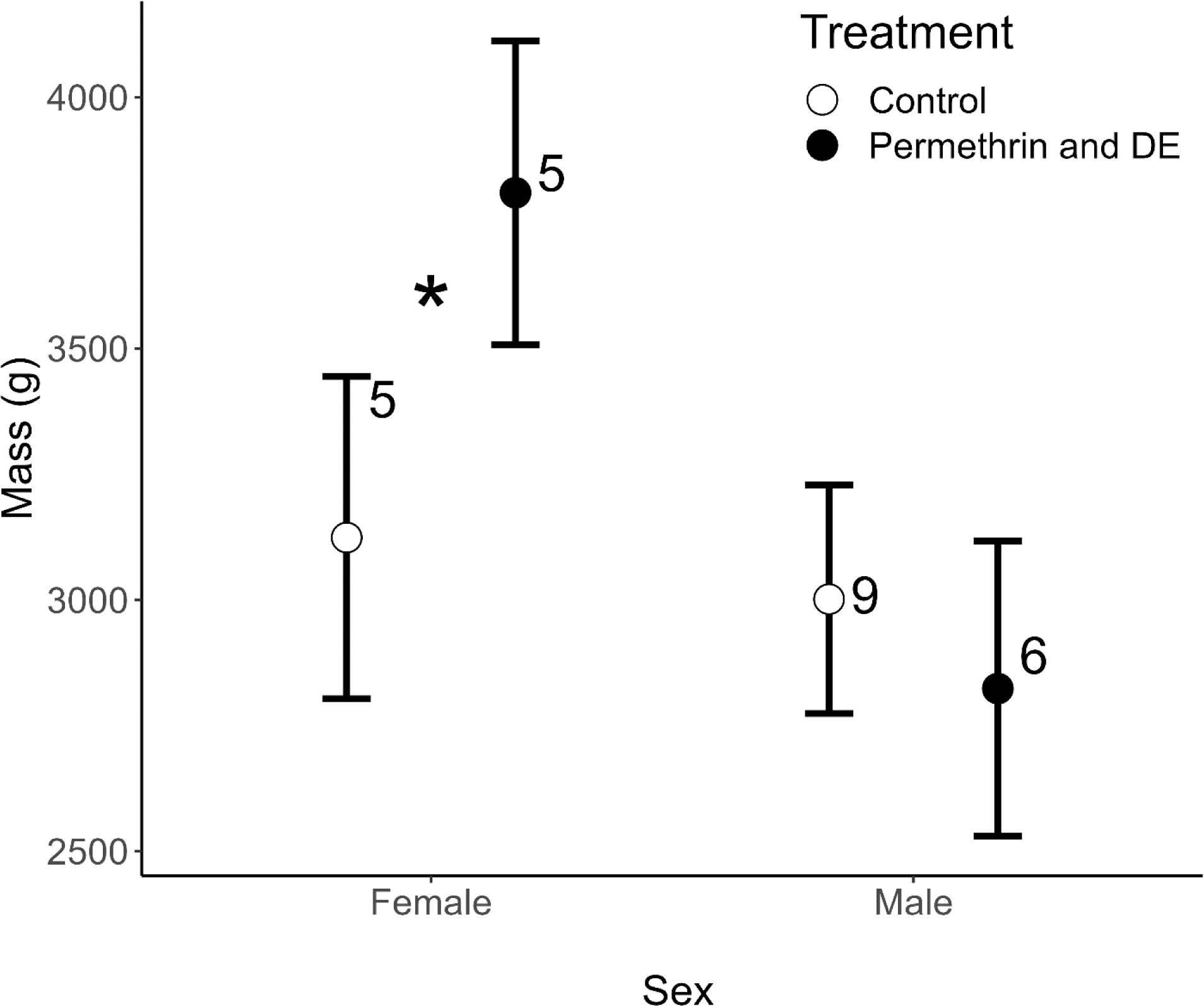
The interaction between sex and parasite treatments on the body mass of golden eagle nestlings > 5 weeks old. Females had significantly higher body mass in permethrin and diatomaceous earth treated nests compared to controls, but there was no effect of parasite treatment on male body mass. Numbers represent sample sizes; circles represent means and bars are 90% confidence limits.

**Figure 5.**
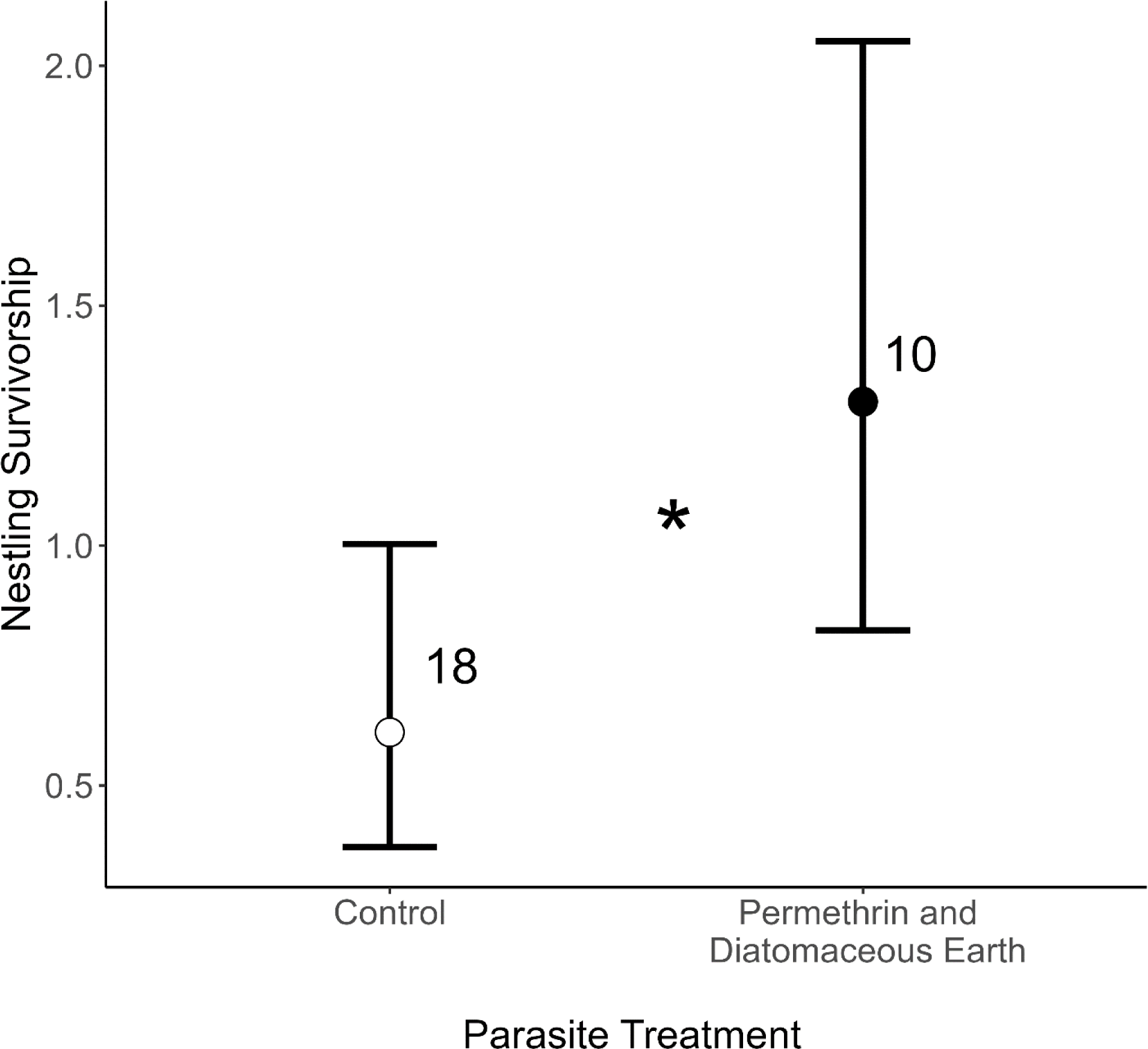
The effect of parasite treatment on golden eagle nestling survivorship. More nestlings successfully fledged from nests that had been treated with permethrin and diatomaceous earth compared with control nests. Numbers represent sample sizes; circles represent means and bars are 90% confidence limits.

### Population Effects

We estimated λ with no trichomonosis treatment to be 1.14 (90% CI: 1.09— 1.19) and with trichomonosis treatment to be 1.18 (90% CI: 1.13—1.24), a difference of 0.04 (90% CI: −0.01— 0.10; Figure 6). The number of adult eagles potentially added from trichomonosis treatments was 0.23 (90% CI: 0.13 — 0.36) per treated territory and a total of 6.13 (90% CI: 3.45 — 9.90) within the 27 territories checked for trichomonosis.

**Figure 6.**
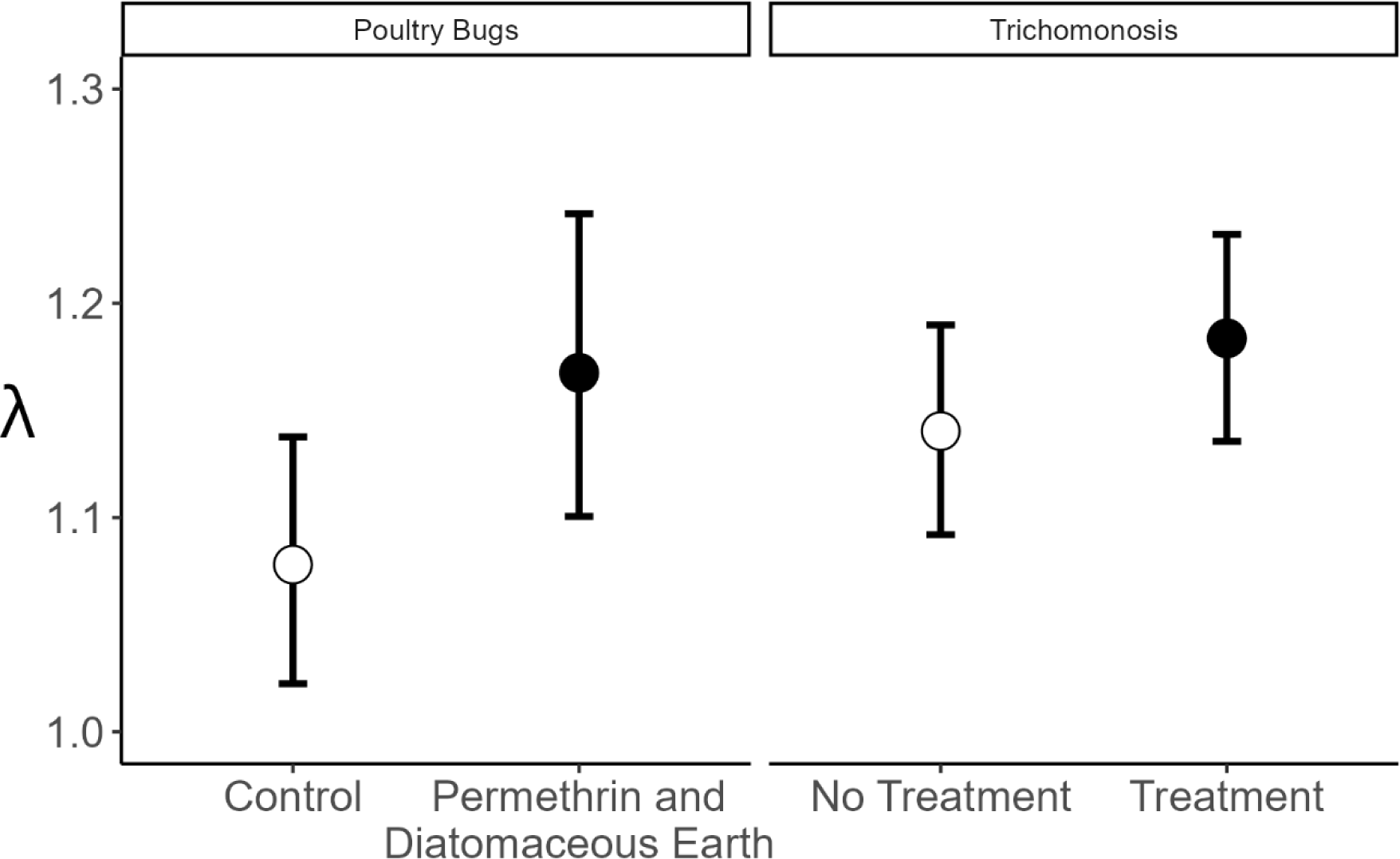
Mean (points), 5^th^ and 95^th^ quantiles (bars) of simulated distributions of the population growth rate (λ) for treatment and control nests for trichomonosis and permethrin and diatomaceous earth treatment and control nests for poultry bugs.

We estimated λ with no parasite treatment to be 1.08 (90% CI: 1.02 — 1.14) and with permethrin and diatomaceous earth treatments to be 1.17 (90% CI: 1.10 — 1.24), a difference of 0.09 (90% CI: 0.01 — 0.17; Figure 6). The number of additional adult eagles resulting from parasite treatments was 0.41 (90% CI: 0.12 — 0.70) per treated territory and a total of 4.09 (90% CI: 1.18 — 7.08) for the 10 treated territories with nesting eagles (Figure 7). In sum, we estimate saving 17.51 (90% CI: 10.50— 25.47) nestlings which equates to 10.24 (90% CI: 6.12— 15.04) adult eagles through disease and parasite treatments.

**Figure 7.**
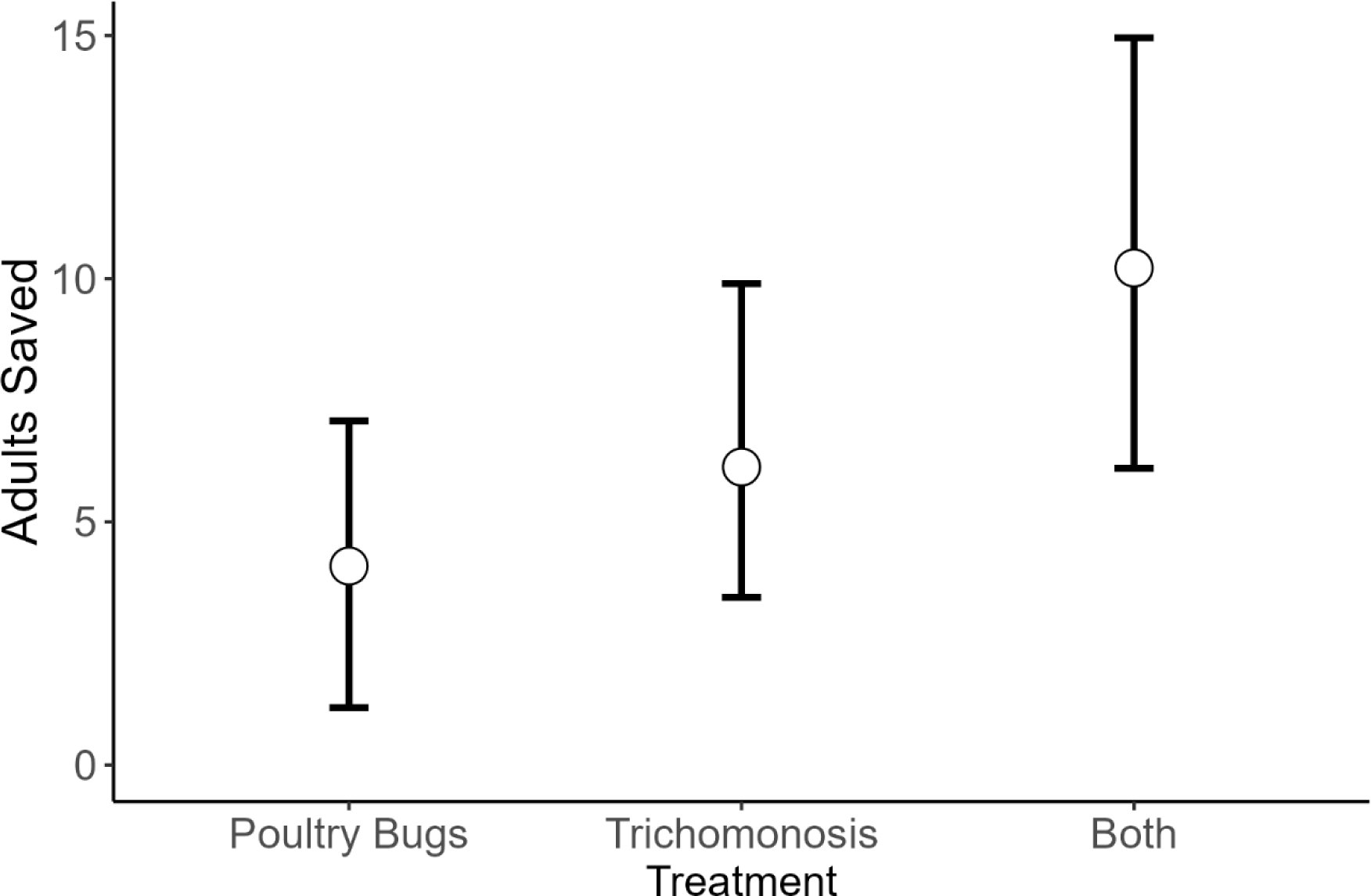
Estimated number adult eagles saved as a result of trichomonosis treatment, poultry bug treatments, and both combined. Points represent the mean (points), and bars the 5^th^ and 95^th^ quantiles of simulated distributions.

## Discussion

Trichomonosis and poultry bugs were a substantial threat to golden eagle nestling survivorship. We successfully treated most nestlings infected with trichomonosis with one dose of an anti-protozoan drug. The permethrin and diatomaceous earth treatment significantly reduced parasite loads in nests and these reductions improved nestling health. Specifically, nestlings were less anemic, female nestlings had higher mass, and nestling survivorship was higher. The parasite treatment, combined with disease treatments, resulted in 17.5 golden eagle nestlings surviving to fledge that otherwise would have died. This equates to an additional 10.2 adult eagles in the population. Given the high certainty of threat locations, treatment success, and the quantifiable nature of treatment, managing disease and parasite threats for golden eagles is an excellent mitigation option.

We found that the majority of nestling deaths (76%) were attributable to disease and parasites. This level of mortality significantly reduced fecundity and had consequences for population growth (see below). The prevalence of trichomonosis among eagle nestlings was high, but this disease was highly treatable with just one dose of an anti-protozoan drug if we caught it early, before plaques were large. A high rate of trichomonosis can be attributed to eagle consumption of rock pigeons, in the absence of black-tailed jackrabbits, their preferred prey, or other suitable prey (Heath et al., 2021); underscoring the ecological repercussions of declining prey populations and trophic cascades. The elimination of this threat would require restoring shrub systems and associated herbivores, like jackrabbits, or removing rock pigeons so they are no longer a vector. Both of these options are challenging, protracted, and expensive. Dependence on alternative food sources, while indicative of the eagles’ adaptability, could pose a significant conservation challenge. If regional reliance on rock pigeons persists, it could potentially lead to the establishment of a conservation-reliant population, wherein ongoing human intervention is required to sustain their reproductive success. Poultry bugs are similarly challenging to manage because these parasites are not host-specific. Nests of other bird species on the same cliffs or rocky outcrops as eagles may become infested and may act as vectors or reservoirs for the poultry bugs (McFadzen and Marzluff 1996). Therefore, the management of disease and parasite threats will likely require repeatable, long-term strategies.

We significantly decreased poultry bug loads in eagle territories with permethrin and diatomaceous earth, which had significant, positive effects on eagle nestling health and survival. Poultry bug loads in eagle nests have increased in recent years likely because of higher overwinter survival of poultry bugs with warming winters (Brown and Brown, 1986). We reduced numbers of poultry bugs before overwintering by applying treatments after eagles left the nest. Post-breeding treatment decreased the chance of long-term side effects on nestlings from permethrin (Bulgarella et al., 2020) because Permacap is estimated to last 90 days and is unlikely to affect eagles in subsequent seasons. However, how the application affects other arthropods in the nests or other animals that use eagle nest structures (e.g., other birds and snakes) should be considered.

There are several remaining questions to address to fine-tune the treatment of poultry bugs in eagle nests. We applied permethrin with diatomaceous earth to increase insect susceptibility to the neurotoxin, because diatomaceous earth scratches exoskeletons. However, it is unclear if the combination is needed or whether permethrin could be effectively applied alone. Diatomaceous earth applied on its own did not affect poultry bug load. Unlike permethrin, which degrades, diatomaceous earth can persist in nests long term. Future research should compare the efficacy of permethrin on its own with permethrin combined with diatomaceous earth. Further, given that poultry bugs can live in other species’ nests and disperse across nest substrates (Heath unpub dat), it is unclear how often territories need to be treated to reduce poultry bug loads. Research to investigate how often territories should be treated is important. Finally, rappelling into eagle nests brings potential harm to the nest structures and the crew. Future work to use uncrewed aerial vehicles to provide remote applications may be a safe and effective alternative.

Eagle health improved after parasite removal. Given the strong effects of poultry bugs on eagle health (Dudek et al., 2021), this result is not surprising. Lower prey quality in recent years (Heath et al., 2021) might make it more difficult for eagles to compensate for the energetic demands of coping with parasites. Nestlings in treated nests had higher hematocrit, were less anemic, and female nestlings were heavier compared to nestlings from control nests. It was interesting that only female mass increased after treatment and not males. However, this result is consistent with other studies of parasite removal that found sex-specific responses (Wolf et al., 2023). Female eagles are larger than male eagles and must grow larger than males during the nestling stage. Therefore, females may have a stronger response to the relaxed pressure of parasitism. Indeed, we found sex-specific responses of eagle nestlings to parasitism in previous work. Specifically, female telomeres but not male telomeres were shorter in parasitized nests (Dudek et al., 2021).

In addition to improved health, nestling eagles had higher survivorship in permethrin and diatomaceous earth-treated nests. This effect is probably because nestlings with high parasite loads jump early from the nest and die, or they die from anemia and starvation (Dudek et al., 2021). A recent paper by Murphy et al. (2023) showed that even if nestling eagles survived parasitized nests long enough to fledge, many of them die in their first year, presumably because they are in poor condition. Our unpublished data tracking juvenile eagles supports this hypothesis. Consequently, the positive effect of parasite treatment on eagle survivorship is likely to be even greater than what we documented by studying survival to fledging, and not longer.

Disease and parasite treatments had projected population-level consequences for golden eagles. This result was surprising because the population viability of long-lived birds, like golden eagles, is most sensitive to changes in adult survival. However, in this population, parasites and disease caused 76% of the deaths of nestlings, and a high proportion (39%) of all nestlings died. Projected population growth rates increased by 4% from trichomonosis treatment and 8% from poultry bug treatment. The projected disease and parasite effects on λ were greater than the estimated population-level effect of lead (0.8%, Slabe et al., 2022), a commonly cited threat to eagles. In combination, poultry bug parasite and trichomonosis disease treatments resulted in 17.5 golden eagle nestlings surviving to fledge over two years that likely would have died without this direct management intervention. Combined, these interventions could be used to offset the take of 10.2 adult eagles.

The expansion of alternative energy infrastructure, habitat fragmentation, declining prey bases, lead exposure, and vehicle collisions threaten eagles across their range and cause additive mortality within populations (e.g., Heath et al., 2021; Katzner et al., 2012; Knick and Dyer 1997, Murphy et al. 2023). Determining how to counteract these threats by off-setting eagle take is a high priority. Further, parasites and disease threats are likely to worsen with climate change and habitat degradation. Treating parasites and disease is an effective compensatory mitigation option because it has population-level effects, while being reliable, cost-effective, and quantifiable. Further, disease and parasite treatments impact populations faster than other management options (i.e., habitat restoration). Given the high effectiveness of disease and parasite treatments, and the easily documented positive effect that these treatments have on eagle nestling survival, combined disease and parasite treatment for golden eagles seems like an excellent option for management to increase population size and to offset take of adult eagles resulting from the expansion of renewable energy infrastructure in the western United States.

## Acknowledgments

Financial and logistical support was provided by the Bureau of Land Management (Award # L14AC00342). The Hycroft Mining Corporation provided funding for this work. We thank H. Beeler for supporting this research and management concept. We thank our partners M. Kochert, J. Weldon, K. Warner, M. Stuber, K. Steenhof, B. Woodbridge, B. Millsap, R. Murphy, B. Pendleton, and I. Robertson, who contributed substantially to this work. M. Henderson, E. Arnold, T. Ely, S. Alsup, J. Wilson, N. Honkomp, C. Pozzangra, J. Harrison, S. Crane, C. Rankin, L. Echavez, and S. Scott conducted fieldwork, placed cameras for monitoring, and helped develop this project. J. Taylor, C. Granthon, and C. Lott provided useful feedback that improved the manuscript.

## Author Contributions

JH and CD conceived of the experiment and wrote the proposal. CD, BD, KM, ER, AS developed methods, collected and managed data. JH, CD, CM, and BR analyzed data with contributions from all co-authors. JH wrote the manuscript with contributions from all co-authors.

## References

Altizer, S., R. Bartel, and B. A. Han. 2011. "Animal Migration and Infectious Disease Risk." Science 331: 296–302.

Allison, T. D., J. F. Cochrane, E. Lonsdorf, and C. Sanders-Reed. 2017. “A Review of Options for Mitigating Take of Golden Eagles at Wind Energy Facilities.” Journal of Raptor Research 51: 319–333

Bale, J. S., and S. A. L. Hayward. 2010. "Insect Overwintering in a Changing Climate." Journal of Experimental Biology 213: 980–994.

Bennett, D. C., A. Yee, Y. J. Rhee, and K. M. Cheng. 2011. “Effect of Diatomaceous Earth on Parasite Load, Egg Production, and Egg Quality of Free-Range Organic Laying Hens.” Poultry Science 90:1416–1426.

Blanco, G., Ó. P. Frías, M. Aida Carrete. 2022. "Oral Disease is Linked to Low Nestling Condition and Brood Size in a Raptor Species Living in a Highly Modified Environment." Current Zoology 69: 109–120.

van den Brand, J. M. A., O. Krone, P. U. Wolf, M. W. G. van de Bildt, G. van Amerongen, A. D. M. E. Osterhaus, and T. Kuiken. 2015. "Host-specific Exposure and Fatal Neurologic Disease in Wild Raptors from Highly Pathogenic Avian Influenza Virus H5N1 During the 2006 Outbreak in Germany." Veterinary Research 46: 24.

Brown, C. R., and M. B. Brown. 1986. “Ectoparasitism as a Cost of Coloniality in Cliff Swallows (*Hirundo pyrrhonota*).” Ecology 67: 1206–1218.

Brown, C. R., S. L. Hannebaum, A. Eaton-Clark, W. Booth, and V. A. O’Brien. 2022. "Elevated Temperature Reduces Overwintering Survival of an Avian Ectoparasite, the Swallow Bug (Hemiptera: Cimicidae: *Cimex vicarius*)." Environmental Entomology 51: 513–520.

Bulgarella, M., S. A. Knutie, M. A. Voss, F. Cunninghame, B. J. Florence-Bennett, G. Robson, R. A. Keyzers, L. M. Taylor, P. J. Lester, G. E. Heimpel, and C. E. Causton. 2020. “Sub-lethal Effects of Permethrin Exposure on a Passerine: Implications for Managing Ectoparasites in Wild Bird Nests.” Conservation Physiology 8: coaa076.

Dawson, R. D. 2004. “Efficacy of Diatomaceous Earth at Reducing Populations of Nest-dwelling Ectoparasites in Tree Swallows.” Journal of Field Ornithology 75: 232–238.

Driscoll, D. E. 2010. Protocol for Golden Eagle Occupancy, Reproduction, and Prey Population Assessment. American Eagle Research Institute, Apache Junction, Arizona, USA, p. 55.

Dudek, B. M., M. T. Henderson, S. F. Hudon, E. J. Hayden, and J. A. Heath. 2021. “Hematophagous Ectoparasites Lower Survival of and Have Detrimental Physiological Effects on Golden Eagle Nestlings.” Conservation Physiology 9: coab060.

Dudek, B. M., M. N. Kochert, J. G. Barnes, P. H. Bloom, J. M. Papp, R. W. Gerhold, K. E. Purple, K. V. Jacobson, C. R. Preston, C. R. Vennum, J. W. Watson, and J. A. Heath. 2018. “Prevalence and Risk Factors of *Trichomonas gallinae* and Trichomonosis in Golden Eagles (*Aquila chrysaetos*) Nestlings in Western North America.” Journal of Wildlife Diseases 54: 755–764.

Fessl, B., S. Kleindorfer, S., and S. Tebbich. 2006. “An Experimental Study on the Effects of an Introduced Parasite in Darwin’s finches.” Biological Conservation 127: 55–61.

Grubb, T. G., W. L. Eakle, and B. N. Tuggle. 1986. “*Haematosiphon inodorus* (Hemiptera: Cimicidae) in a Nest of a Bald Eagle (*Haliaetus leucocephalus*) in Arizona.” Journal of Wildlife Disease 22: 125–27.

Hanssen, S. A., J. O. Bustnes, L. Schnug, S. Bourgeon, T. V. Johnsen, M. Ballesteros, C. Sonne, D. Herzke, I. Eulaers, V. L. B. Jaspers, A. Covaci, M. Eens, D. J. Halley, T. Moum, R. A. Ims, and K. E. Erikstad. 2013. “Antiparasite Treatments Reduce Humoral Immunity and Impact Oxidative Status in Raptor Nestlings.” Ecology and Evolution 3: 5157–5166.

Harrison, J. T., M. N. Kochert, B. P. Pauli, and J. A. Heath. 2019. "Using Motion-Activated Trail Cameras to Study Diet and Productivity of Cliff-Nesting Golden Eagles." Journal of Raptor Research 53: 26–37.

Hassell, J. M., M. Begon, M. J. Ward, and E. M. Fèvre. 2017. "Urbanization and Disease Emergence: Dynamics at the Wildlife–Livestock–Human Interface." Trends in Ecology & Evolution 32: 55–67.

Harvell, C. D., C. E. Mitchell, J. R. Ward, S. Altizer, A. P. Dobson, R. S. Ostfeld, and M. D. Samuel. 2002. “Climate Warming and Disease Risks for Terrestrial and Marine Biota.” Science 296: 2158–62.

Heath, J. A., M. N. Kochert, and K. Steenhof. 2021. “Golden Eagle Dietary Shifts Following Wildfire and Shrub Loss have Negative Consequences for Nestling Survivorship.” Ornithological Applications 123: duab034.

Hickman, G. L. 1968. “The Ecology and Breeding Biology of the Golden Eagle in Southwestern Idaho and Southeastern Oregon.” U.S. Department of the Interior, Division of Wildlife Services.

Hund, A. K., J. T. Blair, and F. W. Hund. 2015. “A Review of Available Methods and Description of a New Method for Eliminating Ectoparasites from Bird Nests.” Journal of Field Ornithology 86: 191–204.

Katzner, T. E., M. N. Kochert, K. Steenhof, C. L. McIntyre, E. H. Craig, and T. A. Miller. 2020. “Golden Eagle (*Aquila chrysaetos*)” version 2.0. In Birds of the World (P. G. Rodewald and B. K. Keeney, Editors). Cornell Lab of Ornithology, Ithaca, NY, USA.

Katzner, T., B., W. Smith, T. A. Miller, D. Brandes, J. Cooper, M. Lanzone, D. Brauning, C. Farmer, S. Harding, D. E. Kramar, C. Koppie, C. Maisonneuve, M. Martell, E. K. Mojica, C. Todd, J. A. Tremblay, M. Wheeler, D. F. Brinker, T. E. Chubbs, R. Gubler, K. O’Malley, S. Mehus, B. Porter, R. P. Brooks, B. D. Watts, and K. L. Bildstein. 2012. “Status, Biology, and Conservation Priorities for North America’s Eastern Golden Eagle (*Aquila chrysaetos*) Population.” The Auk 129: 168–176.

Knick, S.T., and D. L., Dyer. 1997. “Distribution of Black-tailed Jackrabbit Habitat Determined by GIS in Southwestern Idaho.” Journal of Wildlife Management 61: 75–85.

Kochert, M. N., and K. Steenhof. 2012. “Frequency of Nest Use by Golden Eagles in Southwestern Idaho.” Journal of Raptor Research 46: 239–247.

Korunić, Z. 2013. “Diatomaceous Earth - Natural Insecticides.” Pesticidi i Fitomedicina 28: 77–95.

Lafferty, K. D. 2009. "The Ecology of Climate Change and Infectious Diseases." Ecology 90: 888–900.

Larsen, E., J. M. Azerrad, and N. Nordstrom. 2004. Management Recommendations for Washington’s Priority Species, Volume IV: Birds. Washington Department of Fish and Wildlife, Olympia, Washington.

McClure, C. J. W., J. R. S. Westrip, J. A. Johnson, S. E. Schulwitz, M. Z. Virani, R. Davies, A. Symes, H. Wheatley, R. Thorstrom, A. Amar, R. Buij, V. R. Jones, N. P. Williams, E. R. Buechley, and S. H. M. Butchart. (2018). "State of the World’s Raptors: Distributions, Threats, and Conservation Recommendations." Biological Conservation 227: 390–402.

López-Arrabé, J., A. Cantarero, L. Pérez-Rodríguez, A. Palma, and J. Moreno. 2014. “Experimental Pyrethroid Treatment Underestimates the Effects of Ectoparasites in Cavity-nesting Birds due to Toxicity.” Ibis 156: 606–614.

McFadzen, M. E., M. S. Vekasy, T. Y. Morishita, and J. H. Greve. 1996. “Northern Range Extension for *Haematosiphon Inodorus* (Duges) (Hemiptera: Cimicidae).” Pan-Pac Entomologist 72: 41–42.

McFadzen, M. E., and J. M. Marzluff. 1996. “Mortality of Prairie Falcons During the Fledging-Dependence Period.” The Condor 98: 791–800.

Millsap, B. A., T. G. Grubb, R. K. Murphy, T. Swem, and J. W. Watson. 2015. "Conservation Significance of Alternative Nests of Golden Eagles." Global Ecology and Conservation 3: 234–241.

Millsap, B. A., G. S. Zimmerman, W. L. Kendall, J. G. Barnes, M. A. Braham, B. E. Bedrosian, D. A. Bell, P. H. Bloom, R. H. Crandall, R. Domenech, D. Driscoll, A. E. Duerr, R. Gerhardt, S. E. J. Gibbs, A. R. Harmata, K. Jacobson, T. E. Katzner, R. N. Knight, M. J. Lockhart, C. McIntyre, R. K. Murphy, S. J. Slater, B. W. Smith, J. P. Smith, D. W. Stahlecker, and J. W. Watson (2022). "Age-specific Survival Rates, Causes of Death, and Allowable Take of Golden Eagles in the Western United States." Ecological Applications 32: e2544.

Møller, A. P., S. Merino, J. J. Soler, A. Antonov, E. P. Badás, M. A. Calero-Torralbo, F. de Lope, T. Eeva, J. Figuerola, E. Flensted-Jensen, L. Z. Garamszegi, S. González-Braojos, H. Gwinner, S. A. Hanssen, D. Heylen, P. Ilmonen, K. Klarborg, E. Korpimäki, J. Martínez, J. Martínez-de la Puente, A. Marzal, E. Matthysen, P. Matyjasiak, M. Molina-Morales, J. Moreno, T. A. Mousseau, J. T. Nielsen, P. L. Pap, J. Rivero-de Aguilar, P. Shurulinkov, T. Slagsvold, T. Szép, E. Szöllősi, J. Török, R. Vaclav, F. Valera, and N. Ziane. 2013. “Assessing the Effects of Climate on Host-Parasite Interactions: A Comparative Study of European Birds and Their Parasites.” PLoS ONE 8: e82886.

Murphy, R. K., B. A. Millsap, D. W. Stahlecker, C. W. Boal, B. W. Smith, S. D. Mullican, and C. C. Borgman. 2023. “Ectoparasitism and Energy Infrastructure Limit Survival of Preadult Golden Eagles in the Southern Great Plains.” Journal of Raptor Research 57: 505–521.

Palma, L., P. Beja, M. Pais, and L. C. da Fonseca. 2006. “Why do Raptors Take Domestic Prey? The Case of Bonelli’s Eagles and Pigeons.” Journal of Applied Ecology 43: 1075–1086.

Philips, J. R., and D. L. Dindal. 1977. “Raptor Nests as a Habitat for Invertebrates: A Review.” Journal of Raptor Research 11: 87–96.

Philips, J. R. 2007. Pathology: Ectoparasites. – In Bird, D. M. and Bildstein, K. L. (eds.), Raptor Research and Management Techniques. Hancock House, pp. 311–317.

Real, J., S. Manosa, and E. Munoz. 2000. “Trichomoniasis in a Bonelli’s Eagle Population in Spain.” Journal of Wildlife Disease 36: 64–70.

Schaub, M., and M. Kéry. 2022. Integrated Population Models. Elsevier Academic Press, London, UK.

Slabe, V. A., J. T. Anderson, B. A. Millsap, J. L. Cooper, A. R. Harmata, M. Restani, R. H. Crandall, B. Bodenstein, P. H. Bloom, T. Booms, J. Buchweitz, R. Culver, K. Dickerson, R. Domenech, E. Dominguez-Villegas, D. Driscoll, B. W. Smith, M. J. Lockhart, D. McRuer, T. A. Miller, P. A. Ortiz, K. Rogers, M. Schwarz, N. Turley, B. Woodbridge, M. E. Finkelstein, C. A. Triana, C. R. DeSorbo, and T. E. Katzner. 2022. "Demographic Implications of Lead Poisoning for Eagles Across North America." Science 375: 779–782.

Smith, K. F., D. F. Sax, and K. D. Lafferty. 2006. “Evidence for the role of infectious disease in species extinction and endangerment.” Conservation Biology 20: 1349–1357.

Soderlund, D. M., J. M. Clark, L. P. Sheets, L. S. Mullin, V. J. Piccirillo, D. Sargent, J. T. Stevens, and M. L. Weiner. 2002. “Mechanisms of Pyrethroid Neurotoxicity: Implications for Cumulative Risk Assessment.” Toxicology 171: 3–59.

Steenhof, K., M. N. Kochert, and T. L. McDonald. 1997. "Interactive Effects of Prey and Weather on Golden Eagle Reproduction." Journal of Animal Ecology 66: 350–362.

U.S. Department of the Interior. 1979. Snake River Birds of Prey Special Research Report to the Secretary of the Interior. U.S.D.I. Bureau of Land Management, Boise District, Boise, ID, USA.

Vidaña, B., N. Busquets, S. Napp, E. Pérez-Ramírez, M. Á. Jiménez-Clavero, and N. Johnson. 2020. "The Role of Birds of Prey in West Nile Virus Epidemiology." Vaccines 8: 550.

Wiens, J. D., P. S. Kolar, W. G. Hunt, T. Hunt, M. R. Fuller, and D. A. Bell. 2018. “Spatial Patterns in Occupancy and Reproduction of Golden Eagles During Drought: Prospects for Conservation in Changing Environments.” The Condor 120: 106–124.

Wilson, N., and G. Oliver. 1978. “Noteworthy Records of Two Ectoparasites (Cimididae and Hippoboscidae) from the Turkey Vulture in Texas.” The Southwestern Naturalist 23: 305–307.

Wolf, S. E., S. Zhang, and E. D. Clotfelter. 2023. “Experimental Ectoparasite Removal has a Sex-specific Effect on Nestling Telomere Length.” Ecology and Evolution 13: e9861.

